# Using Structural Equation Modeling to Jointly Estimate Maternal and Foetal Effects on Birthweight in the UK Biobank

**DOI:** 10.1101/160044

**Authors:** Nicole M Warrington, Rachel Freathy, Michael C. Neale, David M Evans

## Abstract

**Background:** To date, 60 genetic variants have been robustly associated with birthweight. It is unclear whether these associations represent the effect of an individual’s own genotype on their birthweight, their mother’s genotype, or both.

**Methods:** We demonstrate how structural equation modelling (SEM) can be used to estimate both maternal and foetal effects when phenotype information is present for individuals in two generations and genotype information is available on the older individual. We conduct an extensive simulation study to assess the bias, power and type 1 error rates of the SEM and also apply the SEM to birthweight data in the UK Biobank study.

**Results:** Unlike simple regression models, our approach is unbiased when there is both a maternal and foetal effect. The method can be used when either the individual’s own phenotype or the phenotype of their offspring is not available, and allows the inclusion of summary statistics from additional cohorts where raw data cannot be shared. We show that the type 1 error rate of the method is appropriate, there is substantial statistical power to detect a genetic variant that has a moderate effect on the phenotype, and reasonable power to detect whether it is a foetal and/or maternal effect. We also identify a subset of birth weight associated SNPs that have opposing maternal and foetal effects in the UK Biobank.

**Conclusions:** Our results show that SEM can be used to estimate parameters that would be difficult to quantify using simple statistical methods alone.

**Key Messages:**

- We describe a structural equation model to estimate both maternal and foetal effects when phenotype information is present for individuals in two generations and genotype information is available on the older individual.
- Using simulation, we show that our approach is unbiased when there is both a maternal and foetal effect, unlike simple linear regression models. Additionally, we illustrate that the structural equation model is largely robust to measurement error and missing data for either the individual’s own phenotype or the phenotype of their offspring.
- We describe how the flexibility of the structural equation modelling framework will allow the inclusion of summary statistics from studies that are unable to share raw data.
- Using the structural equation model to estimate the maternal and foetal effects of known birthweight associated loci in the UK Biobank, we identify three loci that have primary effects through the maternal genome and six loci that have opposite effects in the maternal and foetal genomes.

## Introduction

Birthweight is a complex trait and low birthweight is robustly associated with increased risk of a range of cardio-metabolic diseases in later life(1). It has long been known that birthweight is under the influence of both maternal and foetal genetic sources of variation. Using a large sample consisting of the offspring of twins, Magnus illustrated that more than 50% of the variation in birthweight is caused by fetal genes and less than 20% was caused by maternal genes(2). Subsequent studies have reported lower proportions of the variance explained by both fetal and maternal genes, but all have shown that the fetal contribution is larger than the maternal contribution(3, 4). Using a method that partitions trait variance into components due to the maternal and foetal genomes(5), we reported that common genetic variants in the fetal genome explained approximately 28% of the variation in birthweight, whereas common genetic variants in the maternal genome only explained approximately 8% of the total variance(6).

We and others have begun to investigate the specific regions of the genome that influence fetal growth using genome-wide association studies (GWAS). In a recent GWAS meta-analysis combining data from the Early Growth Genetics consortium (EGG; http://egg-consortium.org/) and the UK Biobank(7), we identified 60 SNPs associated with birthweight at genome-wide levels of significance(6). One difficulty we faced in interpreting our results was that it was often not clear whether genetic associations reflected the effect of an individual’s own genotype on their birthweight, an effect of their mother’s genotype on their birthweight (i.e., maternal genotype mediated through the intrauterine effect), or some combination of both. For example, rare mutations in the *GCK* gene, which cause a defect in the sensing of glucose by the pancreas, have radically different associations with birthweight according to their parent of origin. If inherited paternally, birthweight is lower due to reduced glucose sensing and consequent reduced insulin secretion, which results in reduced growth. But if maternally inherited (i.e. present in both mother and fetus), birthweight is close to the population average because the maternal hyperglycemia compensates for the fetal defect in glucose sensing. In the case that the mother has hyperglycemia due to a GCK mutation, but the fetus does not inherit the mutation, the birthweight is higher due to normal glucose sensing and thus above-average insulin secretion. This example reflects contrasting effects mediated through the intrauterine environment (i.e. maternal effects) and direct effects of the offspring’s genotype(8).

In an attempt to resolve this question, in Horikoshi *et al*.(6) we first performed a simple linear regression of an individual’s self-reported birthweight on their own genotype, and then for the UK Biobank women, a linear regression of the birthweight of their first born child on their own genotype. We then compared the maternal and foetal effect sizes to get an idea of whether the locus was operating through the maternal or the individual’s own genotype. However, this approach was suboptimal since it did not consider the correlation between maternal and offspring genotypes, and therefore did not accurately estimate the relative importance of these two potential sources of variation. We also examined the genetic associations with birthweight in cohorts that had genotype information on both mother and offspring. Performing an analysis of offspring birthweight on maternal genotype and conditioning on offspring genotype should yield an unbiased estimate of the mother’s genetic influence on her child’s birthweight, and likewise a regression of offspring birthweight on offspring genotype conditioning on maternal genotype should produce an unbiased estimate of the foetal contribution on birthweight. The difficulty however is that there is a paucity of cohorts in the world that have birthweight data as well as genotype data on both mothers and children, meaning that such an analysis is likely to have low power to resolve maternal and foetal effects.

To better estimate the maternal and foetal genetic contributions to birthweight for each of the 60 genome-wide significant variants reported in Horikoshi *et al*.(6) we utilized a structural equation modelling (SEM) approach with birthweight data from the UK Biobank. Our method enables us to model both grand-maternal and offspring genotypes (which were absent in the UK Biobank) as latent factors, and to estimate maternal and foetal effects on birthweight in the same statistical model. To investigate the properties of our approach, we first performed a series of simulations to; i) quantify any bias in the effect estimates for the maternal and foetal effects, and ii) estimate power to detect maternal and foetal effects and type 1 error. We also assessed the effect of allele frequency and measurement error in birthweight (which can often be an issue with self-report) on our estimates. We show that our method provides accurate estimates of maternal and foetal effects under a range of different scenarios and increased power to detect genetic association when maternal and foetal effects operate in opposite directions. We also show how our framework can easily combine summary results data from additional cohorts, including previous large scale GWAS meta-analyses, involving either maternal or offspring phenotypes. Using the UK Biobank data(7), we provide strong evidence to suggest that several of the known birthweight associated SNPs exert effects acting in opposite directions on birthweight through the maternal and foetal genotypes.

## >Methods

### Simulations

We performed simulations to investigate the bias, power and type one error rate of the SEM for modelling both the individual’s own genetic effect (referred to as the “foetal effect”) and maternal genetic effects on birthweight. The model we used for generating the data is illustrated in Figure 1 and the R code used for performing these simulations is provided in the Supplementary Material. For each scenario, we generated 10 000 replicates of 30 000 maternal-offspring pairs. For each replicate we generated grandparental and paternal genotypes at a single locus. Assuming autosomal Mendelian inheritance, additivity and unit variance, latent variables for the genotype of the individual’s mother (i.e., grand-maternal genotype; G_G_), the individual’s own genotype (SNP) and offspring’s genotype (G_o_) were generated. The individual’s own birthweight variable (BW) for each family *i*, was generated using the following equation:

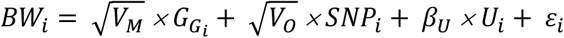

where *V_M_* denotes the variance in birthweight explained by the maternal genotype (“maternal effect”), *G_G_* is a latent variable indexing the genotype of the individual’s mother, *V_O_* is the variance in birthweight explained by the individual’s own genotype (“foetal effect”), *SNP* is the genotype of the individual, *U* is a standard normal random variable representing all residual genetic and environmental sources of similarity between mother and offspring, *ϐ_U_* is the total effect of *U* on the individual’s own birthweight, and *ε* is a random normal variable with mean zero and variance needed to ensure that BW has unit variance asymptotically.

**Figure 1:**
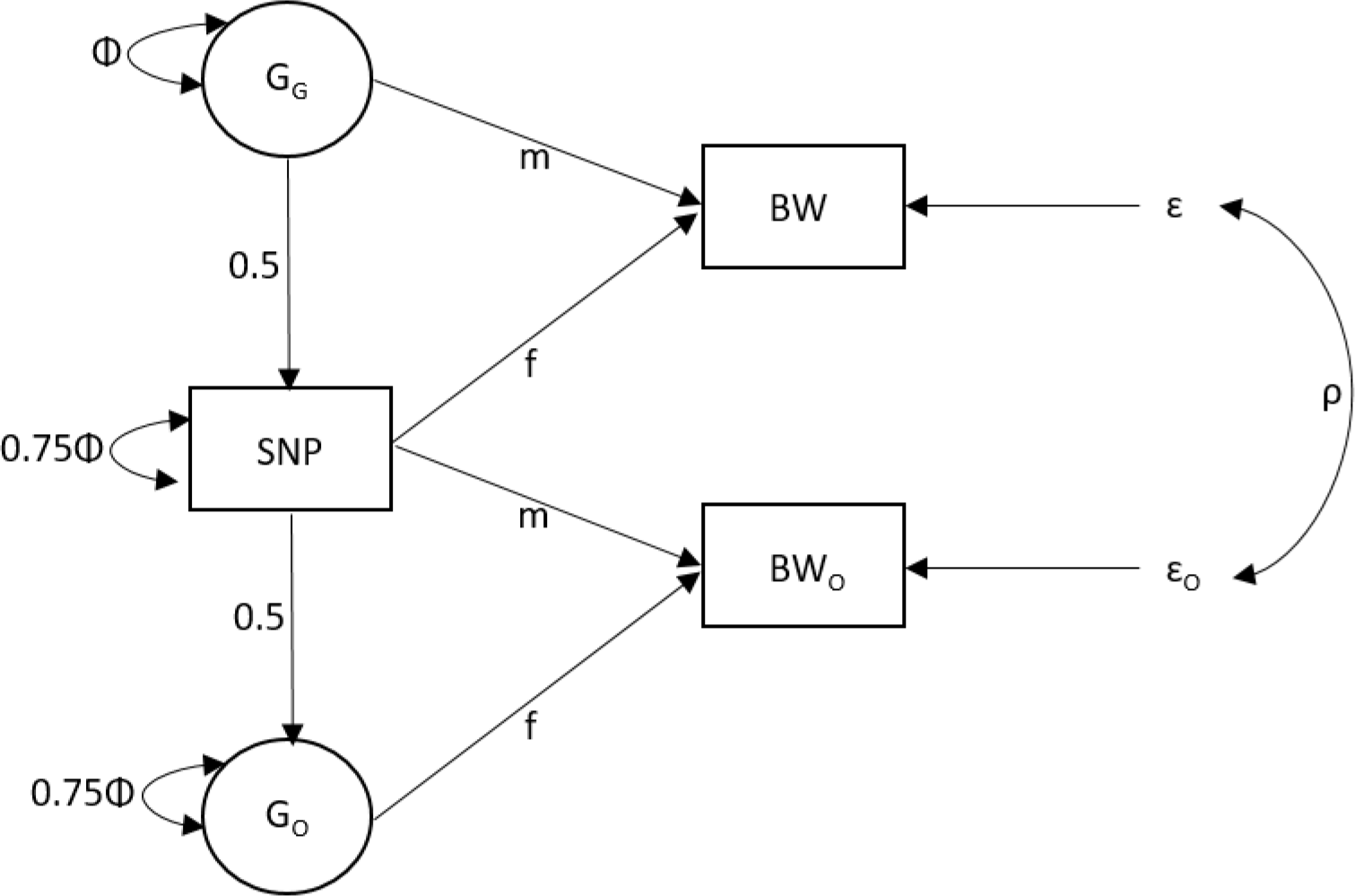
Diagram of the structural equation model (SEM) used for the simulation study and the UK Biobank analysis of birthweight. The three observed variables include the birthweight of the individual (BW), the birthweight of their offspring (BW_o_) and the genotype of the individual (SNP). The variance of the latent genotypes for the individual’s mother (G_G_) and offspring (G_o_) is set to Φ. The m and f path coefficients refer to maternal and foetal effects respectively. The residual error terms for the birthweight of the individual and their offspring are represented by e and eo respectively. The covariance between residual genetic and environmental sources of variation is given by ρ.

Similarly, offspring birthweight (BW_O_) for each family i, was generated using the following equation:

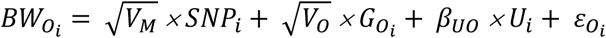

where *G_O_* is a latent variable indexing the offspring genotype, *ϐ_UO_* is the total effect of *U* on offspring birthweight, and *ε_O_* is a random normal variable with mean zero and variance needed to ensure that BW_O_ has unit variance asymptotically.

In all simulations, the regression of phenotype on residual shared genetic and environmental factors was set to 0.5 (i.e., *ϐ_U_*= *ϐ_UO_*= 0.5). We considered the effects of: allele frequency (p = 0.99, p = 0.90 or p = 0.5); the strength of the foetal genetic effect on birthweight (*V_O_*= 0%, *V_O_*= 0.02% or *V_O_*= 0.04%); and the strength of the maternal genetic effect on birthweight (*V_M_* = 0%, *V_M_*= 0.01% or *V_M_*= 0.02%). We simulated the foetal and maternal genetic effects to have both increasing and decreasing effects on birthweight.

For each simulated dataset we fit a series of models:

1. Linear models regressing either the individual’s own birthweight or, for the women, the birthweight of their offspring on the SNP (individual’s own genotype), which respectively estimate the foetal and maternal genetic effects on birthweight. This is equivalent to the model typically used in genetic studies of birthweight(6) and was used for comparison purposes.
2. SEM estimating both maternal and foetal effects as illustrated in Figure 1. P-values were calculated using Wald tests.
3. SEM estimating only the foetal effect. This model was fit to conduct a likelihood ratio test for the maternal effect. This likelihood ratio test P-value was compared to the Wald test P-value for the foetal effect.
4. SEM estimating only the maternal effect to conduct a likelihood ratio test for the foetal effect. This likelihood ratio test P-value were compared to the Wald test P-values for the maternal effect.
5. SEM with neither foetal nor maternal paths (i.e., both fixed to zero). This model was fit to conduct a likelihood ratio test of the overall SNP effect. The P-value from this test is referred to as the two degrees of freedom (2DF) test P-value.

Bias was defined as the mean difference between the estimated SNP effect and the true parameter across the 10 000 simulations and was calculated for both the maternal and foetal effects. A 95% confidence interval was calculated around the bias to give an indication of the uncertainty in the estimate. Power was defined as the proportion of tests that reached P<0.05 under the alternative hypothesis, and type 1 error rate the proportion of tests that reached P<0.05 under the null hypothesis.

### Additional simulations investigating measurement error

In the UK Biobank, female participants were asked to report the birthweight of their first offspring to the nearest pound. After appropriate data cleaning, this left six discrete birthweight values for the offspring (see below). We therefore conducted a second set of simulations to investigate the effect of this type of measurement error, using the same method as described above but rounding the birthweight of the offspring to the nearest unit.

Given that birthweights of both individuals and their offspring are self-reported in the UK Biobank, we also assessed the potential effect of measurement error on both variables. To do this, we added a normally distributed error component to both simulated birthweight measurements, which is referred to as discrimination or classical measurement error(9). For example, an individual’s own birthweight with measurement error was generated as follows:

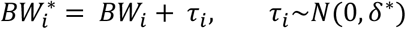

Where δ^*^ was chosen to produce a specific R^2^ value for the regression of BW^*^ on BW, using the following equation:

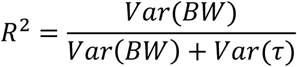

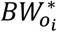 was generated in the same way. We varied the value of R^2^ (1.00, 0.75, 0.50, 0.25), where lower values of R^2^ represent increasing measurement error. We used a subset of maternal and foetal effect sizes to get an idea of whether the impact of measurement error is influenced by effect size; neither a maternal or foetal effect (*V_0_* = 0% = *V_M_*), large foetal effect and no maternal effect (*V_0_* = 0.04%, *V_M_* = 0%), no foetal effect and a large maternal effect (*V_0_* = 0%, *V_M_* = 0.02%), large foetal and maternal effect (*V_0_* = 0.04%, *V_M_* = 0.02%) and large foetal and maternal effect in opposite directions. All simulations were conducted with an allele frequency of p = 0.5.

### Additional simulations investigating missing data

We also assessed the impact of when individuals did not have both their own and their offspring birthweight available. We simulated four additional scenarios, all with minor allele frequency of p = 0.5 and a total sample size of 30 000 individuals; 1) 15 000 individuals with both their own and their offspring’s birthweight and 15 000 individuals with their own birthweight only, 2) 15 000 individuals with both their own and their offspring’s birthweight and 15 000 individuals with their offspring’s birthweight only, 3) 15 000 individuals with both their own and their offspring’s birthweight, 7500 individuals with their own birthweight only and 7500 individuals with their offspring’s birthweight only, 4) 15 000 individuals with their own birthweight only and 15 000 individuals with their offspring’s birthweight only (i.e., no individuals with both birthweight measures and therefore the term *ρ* in Figure 1 could not be estimated). Given that we observed very close correspondence between the likelihood ratio and Wald tests, we only conducted Wald tests because these were computationally easier to perform.

### UK Biobank

UK Biobank phenotype data were available on 502 643 individuals, of which 279 959 reported their own birthweight at either the baseline or follow-up visits. There were 7693 individuals who were part of multiple births and were excluded from the analyses. Of the 9034 individuals who reported their own birthweight at both baseline and follow-up, 401 were excluded because the two values differed by more than 0.5kg. For those individuals who reported different values between baseline and follow-up (<0.5kg) we took the baseline measure for the analyses. Finally, we excluded individuals who reported their own birthweight to be <2.5kg or >4.5kg, as these are implausible for live term births before 1970. In total, 233 662 individuals had data on their own birthweight matching our inclusion criteria.

Women in the UK Biobank were also asked to report the birthweight of their first child to the nearest pound. We used the same inclusion criteria as for their own birthweight leaving 210 405 individuals with birthweight of their first child, 109 205 of whom had also reported their own birthweight.

Genotype data from the May 2015 release were available on a subset of 152 248 individuals. In addition to the quality control metrics performed centrally by the UK Biobank, we excluded individuals who were related. We defined a subset of ‘white European’ ancestry samples using a *K*-means (K=4) clustering approach based on the first four genetically determined principal components. A subset of 89 296 individuals with genotype data, a valid birthweight for themselves or their first child and were genetically of ‘white European’ ancestry were included in the analysis. Of these, 24 962 were men who only reported their own birthweight. Among the women, 8723 reported only their own birthweight, 24 645 reported only that of their first child, and 30 966 reported both. We adjusted both the individual’s own birthweight and the birthweight of their first child for the principal components that were associated with birthweight, adjusted the individual’s own birthweight for sex (sex was not reported for the offspring) and then created z-scores. A subset of 58 autosomal SNPs out of the 60 birthweight associated SNPs(6) were extracted from the imputed files provided by UK Biobank and aligned to the birthweight increasing allele (rs62240962 was not available and rs11096402 is on the X chromosome).

## Results

### Bias

Figure 2 shows the bias calculated from the simulations for the linear model and the SEM in the simulations with allele frequency of 0.5 and all 30 000 individuals whom had complete data for both their own birthweight and the birthweight of their offspring. The foetal effect estimates from the standard linear model are biased wherever there is a maternal effect that is not being modelled. For example, in the scenarios where there is both a foetal and maternal effect, the estimated foetal effect approximately equals the true foetal effect plus half the true maternal effect. In other words, the bias of the estimated foetal effect is approximately half the true maternal effect. In the scenarios where there is no maternal effect, then the foetal effect estimated from the linear model is unbiased. The same pattern of bias occurs for the maternal effect estimates. Conversely, the SEM is unbiased for both the maternal and foetal effects as it simultaneously models both effects. The bias and 95% confidence intervals for all simulation results are presented in Supplementary Table 1.

**Figure 2:**
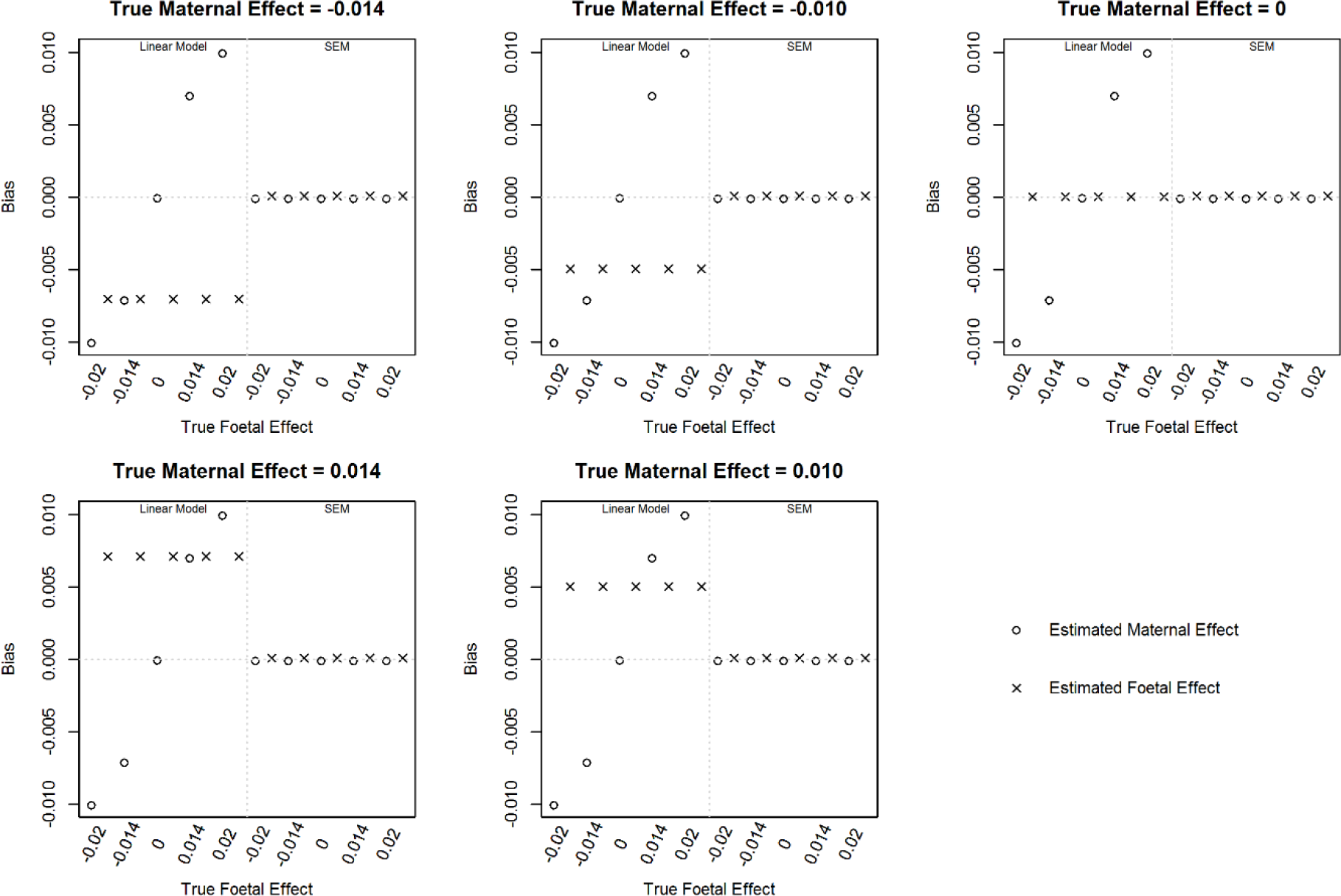
Bias in effect estimates with an allele frequency of 0.5 and varying maternal and foetal effect sizes using two linear models that assess the maternal and foetal effects independently (left panel) or the structural equation model (SEM, right panel) assessing both the maternal and foetal effects simultaneously.

When there was measurement error in either the individual’s own birthweight or the birthweight of the offspring, the estimates of maternal and foetal effects were unbiased (Table 1 for abridged results and full results in Supplementary Table 2). However, there was a decrease in the precision of the estimate (i.e., increase in the standard error) as the measurement error increased (Table 1 for abridged results and full results in Supplementary Table 2). For a small number of scenarios when the birthweight of the offspring was distributed as it is in the UK Biobank, a small bias was introduced from the SEM (Supplementary Table 3); this bias differed across allele frequency and true effect size for both the maternal and foetal effects and no clear pattern was observed. The bias was less than 4% of the true value in all scenarios and substantially lower than the bias introduced in the linear models.

**Table 1:** The effect of measurement error in the individual’s own birthweight and the birthweight of their offspring on bias, precision and power in the structural equation model (SEM).

| Measurement error | | $\sqrt{V_O} = 0; \sqrt{V_M} = 0$ | | | $\sqrt{V_O} = 0; \sqrt{V_M} = -0.014$ | | | $\sqrt{V_O} = -0.020; \sqrt{V_M} = 0$ | | | $\sqrt{V_O} = -0.020; \sqrt{V_M} = -0.014$ | | | $\sqrt{V_O} = 0.020; \sqrt{V_M} = -0.014$ | | |
| --- | --- | --- | --- | --- | --- | --- | --- | --- | --- | --- | --- | --- | --- | --- | --- | --- |
| R<sup>2</sup> BW | R<sup>2</sup> BW<sub>O</sub> | Mean estimate | Mean standard error | Power | Mean estimate | Mean standard error | Power | Mean estimate | Mean standard error | Power | Mean estimate | Mean standard error | Power | Mean estimate | Mean standard error | Power |
| **True Foetal Effect** |  |  |  |  |  |  |  |  |  |  |  |  |  |  |  |  |
| **Measurement error for offspring birthweight only** |  |  |  |  |  |  |  |  |  |  |  |  |  |  |  |  |
| 1.00 | 1.00 | 0.000 | 0.008 | 0.048 | 0.000 | 0.008 | 0.048 | -0.020 | 0.008 | 0.738 | -0.020 | 0.008 | 0.738 | 0.020 | 0.008 | 0.741 |
| 1.00 | 0.75 | 0.000 | 0.008 | 0.050 | 0.000 | 0.008 | 0.049 | -0.020 | 0.008 | 0.711 | -0.020 | 0.008 | 0.712 | 0.020 | 0.008 | 0.702 |
| 1.00 | 0.50 | 0.000 | 0.009 | 0.048 | 0.000 | 0.009 | 0.051 | -0.020 | 0.009 | 0.655 | -0.020 | 0.009 | 0.656 | 0.020 | 0.009 | 0.634 |
| 1.00 | 0.25 | 0.000 | 0.010 | 0.048 | 0.000 | 0.010 | 0.048 | -0.020 | 0.010 | 0.515 | -0.020 | 0.010 | 0.516 | 0.020 | 0.010 | 0.490 |
| 1.00 | UKBB<sup>1</sup> | 0.000 | 0.008 | 0.047 | 0.000 | 0.008 | 0.047 | -0.020 | 0.008 | 0.733 | -0.020 | 0.008 | 0.736 | 0.020 | 0.008 | 0.732 |
| **Measurement error for individuals own birthweight only** |  |  |  |  |  |  |  |  |  |  |  |  |  |  |  |  |
| 1.00 | 1.00 | 0.000 | 0.008 | 0.048 | 0.000 | 0.008 | 0.048 | -0.020 | 0.008 | 0.738 | -0.020 | 0.008 | 0.738 | 0.020 | 0.008 | 0.741 |
| 0.75 | 1.00 | 0.000 | 0.009 | 0.048 | 0.000 | 0.009 | 0.049 | -0.020 | 0.009 | 0.616 | -0.020 | 0.009 | 0.618 | 0.020 | 0.009 | 0.612 |
| 0.50 | 1.00 | 0.000 | 0.011 | 0.047 | 0.000 | 0.011 | 0.047 | -0.020 | 0.011 | 0.448 | -0.020 | 0.011 | 0.449 | 0.020 | 0.011 | 0.453 |
| 0.25 | 1.00 | 0.000 | 0.015 | 0.046 | 0.000 | 0.015 | 0.046 | -0.020 | 0.015 | 0.248 | -0.020 | 0.015 | 0.248 | 0.020 | 0.015 | 0.253 |
| **True Maternal Effect** |  |  |  |  |  |  |  |  |  |  |  |  |  |  |  |  |
| **Measurement error for offspring birthweight only** |  |  |  |  |  |  |  |  |  |  |  |  |  |  |  |  |
| 1.00 | 1.00 | 0.000 | 0.008 | 0.052 | -0.014 | 0.008 | 0.456 | 0.000 | 0.008 | 0.052 | -0.014 | 0.008 | 0.456 | -0.014 | 0.008 | 0.457 |
| 1.00 | 0.75 | 0.000 | 0.009 | 0.047 | -0.014 | 0.009 | 0.350 | 0.000 | 0.009 | 0.047 | -0.014 | 0.009 | 0.351 | -0.014 | 0.009 | 0.351 |
| 1.00 | 0.50 | 0.000 | 0.011 | 0.046 | -0.014 | 0.011 | 0.245 | 0.000 | 0.011 | 0.046 | -0.014 | 0.011 | 0.246 | -0.014 | 0.011 | 0.245 |
| 1.00 | 0.25 | 0.000 | 0.015 | 0.047 | -0.014 | 0.015 | 0.142 | 0.000 | 0.015 | 0.047 | -0.014 | 0.015 | 0.142 | -0.014 | 0.015 | 0.142 |
| 1.00 | UKBB<sup>1</sup> | 0.000 | 0.008 | 0.052 | -0.014 | 0.008 | 0.426 | 0.000 | 0.008 | 0.052 | -0.014 | 0.008 | 0.424 | -0.014 | 0.008 | 0.433 |
| **Measurement error for individuals own birthweight only** |  |  |  |  |  |  |  |  |  |  |  |  |  |  |  |  |
| 1.00 | 1.00 | 0.000 | 0.008 | 0.052 | -0.014 | 0.008 | 0.456 | 0.000 | 0.008 | 0.052 | -0.014 | 0.008 | 0.456 | -0.014 | 0.008 | 0.457 |
| 0.75 | 1.00 | 0.000 | 0.008 | 0.050 | -0.014 | 0.008 | 0.418 | 0.000 | 0.008 | 0.051 | -0.014 | 0.008 | 0.418 | -0.014 | 0.008 | 0.418 |
| 0.50 | 1.00 | 0.000 | 0.009 | 0.051 | -0.014 | 0.009 | 0.375 | 0.000 | 0.009 | 0.051 | -0.014 | 0.009 | 0.374 | -0.014 | 0.009 | 0.375 |
| 0.25 | 1.00 | 0.000 | 0.010 | 0.050 | -0.014 | 0.010 | 0.283 | 0.000 | 0.010 | 0.051 | -0.014 | 0.010 | 0.283 | -0.014 | 0.010 | 0.283 |

<sup>1</sup>UKBB indicates that offspring birthweight distributed as in the UK Biobank, where only 6 discrete values are available

In simulations where either the individual’s own birthweight or that of their offspring was missing, the SEM continued to produce unbiased estimates (Supplementary Table 1). However, in the simulations where all individuals only had their own birthweight or the birthweight of their offspring (i.e., no individuals had both birthweight measures), we detected a small bias in the maternal effect estimate (bias approximately −0.0003, or less than 3% of the true value).

### Power/Type 1 error

The power and type 1 error results for all simulations from the SEM and the linear models for the foetal and maternal effects are presented in Supplementary Table 1.

The linear model has greater power than the Wald test in the SEM when the SNP has either a foetal or maternal effect only (i.e., when the effect estimate is unbiased; Supplementary Table 1). For example, the power is greater for the foetal effect estimated using the linear model over the Wald test from the SEM when the maternal effect is zero. Nevertheless, there is still substantial power to detect an effect using the Wald test in the SEM with α=0.05, with 74% power to detect a variant that explains 0.04% of the variance, 45% power for one explaining 0.02% of the variance, and 25% power for one explaining 0.01% of the variance in a sample of 30 000 individuals with both their own and their offspring’s birthweight. However, the two degree of freedom test has very similar power to the linear model when the SNP had either a foetal or maternal effect only and greater power in most scenarios than testing either maternal or foetal effects individually using the Wald test (Figure 3 and Supplementary Table 1). It is worth nothing that the power estimates from the linear models are artificially inflated due to the bias introduced in the linear models, however we have included them in the figure as they give an indication of what the power of a standard genetic analysis would be. This indicates that the SEM can detect when a SNP affects birthweight but it has lower power to detect whether the effect is driven by the mother or the offspring. For example, when the variant explains 0.04% of the variance using the individual’s own genotype and 0.02% of the variance using the mother’s genotype, with α=0.05 the SEM has 74% power to detect the foetal effect, 45% power to detect the maternal effect and 100% power to detect any effect of the variant using the two degree of freedom test in a sample of 30 000 individuals with both their own and their offspring’s birthweight. Similarly, with ɑ=0.05 and 30 000 individuals with complete data, when the variant explains 0.02% of the variance using the individual’s own genotype and 0.01% of the variance using the mother’s genotype, the SEM has 45% power to detect the foetal effect, 25% power to detect the maternal effect and 95% power to detect any effect of the variant using the two degree of freedom test.

**Figure 3:**
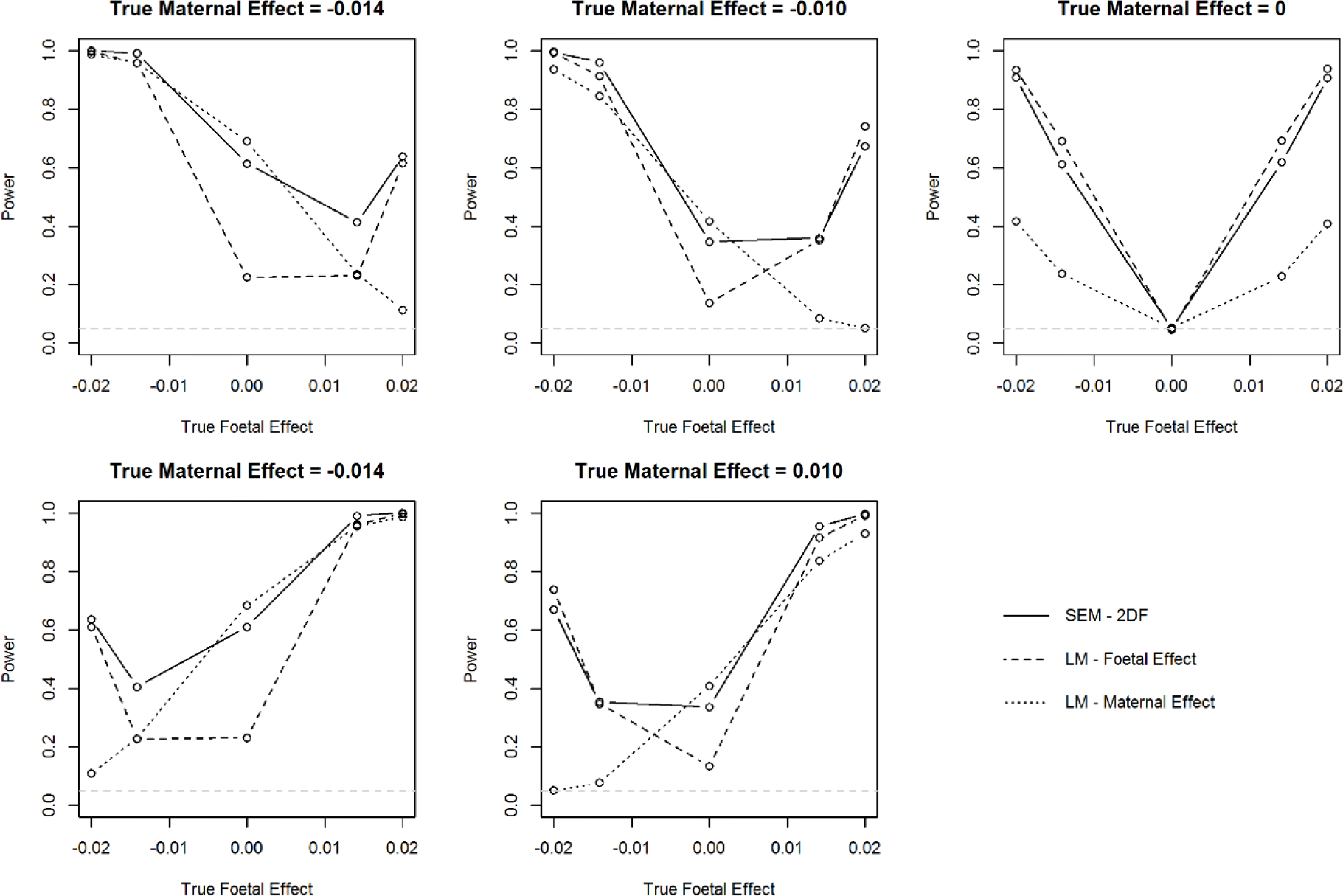
Power of the two degree of freedom test using the structural equation model (SEM) assessing both the maternal and foetal effects simultaneously and power of the two linear models (LM) that assess the maternal and foetal effects independently. Note, power from the linear models is artificially inflated due to the bias in the effect estimate, but they are presented here as a comparison to what would be provided from a standard genetic analysis of birthweight. Power is presented for simulations with a minor allele frequency of 0.5.

Power for both the foetal and maternal effect is reduced when there is measurement error in either the individual’s own birthweight or the birthweight of their offspring due to the decrease in precision of the estimate (Table 1 and Supplementary Table 2). This decrease in power was the same across the different true effect sizes for the foetal and maternal effects.

Figure 4 shows the power when not all individuals have complete data for both maternal and offspring birthweight. Power is greatest when information is available on both the individual’s own and their offspring’s birthweight; however, there is only a small decrease in power to detect the foetal effect when information on the offspring birthweight is not available 50% of the individual’s, and for power to detect the maternal effect when the individual’s own birthweight is not available in 50% of the individual’s. Interestingly, the SEM can still be utilized to estimate maternal and foetal effects when the sample consists of some individuals only measured on their own birthweight, and others who have only reported the birthweight of their offspring. However, the power to detect either a foetal or maternal effect is approximately half of that when birthweight data are available on both individuals in the pair.

**Figure 4:**
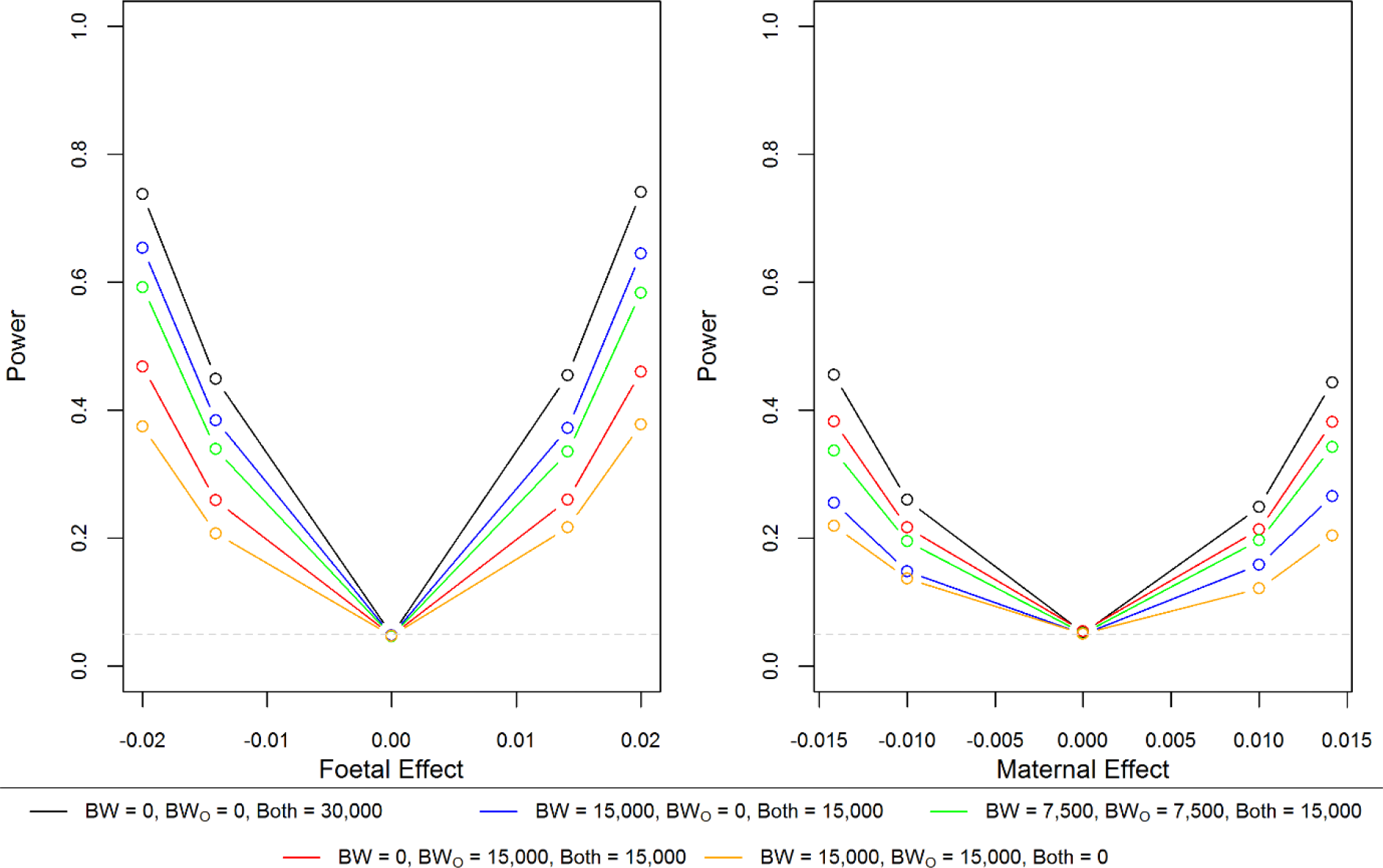
Power from the structural equation model (SEM) with different combinations of individuals reporting their own birthweight (BW) or their offspring’s birthweight (BW_O_). Power for the foetal effect is presented from the simulations where there is no maternal effect; however, similar estimates were obtained when there was a maternal effect (see Supplementary Table 1 for full results). Similarly, for the maternal effect, results are presented from simulations where there is no foetal effect. Power is presented for simulations with a minor allele frequency of 0.5.

As expected, the type 1 error from the linear model is inflated in situations where the estimated effect is biased (Supplementary Table 1). However, the type 1 error is well controlled when using the SEM. It remains controlled when birthweight of the offspring is distributed as in the UK Biobank (Supplementary Table 3), when there is measurement error in either the individuals own birthweight or the birthweight of their offspring (Table 1, Supplementary Table 2), or when data is not available on both the individuals own birthweight and their offspring’s birthweight (Supplementary Table 1).

The difference between P-values estimated using the Wald test and the likelihood ratio test in the SEM was negligible (Supplementary Table 1 for mean difference), indicating that the Wald test was adequate.

### Timing

The SEM can be fitted with either the raw data or observed covariance matrices. As seen in Supplementary Figure 1, the computational time is approximately 100 times faster using the covariance matrices than the raw data. There is not a substantial difference in computational time between datasets with different amounts of missing data for the phenotype of the individual or their offspring when fitting the model using the raw data, but there is a difference for variants with lower minor allele frequencies which take longer to run than common variants. When using covariance matrices, however, it takes slightly longer to fit the model when data are missing for either the individual or their offspring, because the model fits two or three sub-models simultaneously (i.e., one for each of the complete data subsets; one for individuals with both phenotypes, one for individuals with their own phenotype only and one for individuals with their offspring’s phenotype only). The estimates from the model fit with the raw data are the same as those using with covariance matrices. In comparison to a linear model, the SEM using covariance matrices takes about three times as long to compute with a sample size of 30 000 individuals (Supplementary Figure 1).

### UK Biobank

Figure 5 presents the results from the SEM for each of the 58 birthweight associated SNPs in the UK Biobank. It is evident that most of the 58 SNPs only have evidence for a foetal effect, which is unsurprising given how the SNPs were selected. Three SNPs primarily have a maternal effect (*EBF1, ACTL9* and *MTNR1B*) and eight SNPs have evidence for both. Perhaps the most interesting of these six SNPs are those where the birthweight increasing allele identified in Horikoshi *et* al.(6) has a opposite effects on birthweight through the foetal and maternal genotype, half of which are known type 2 diabetes loci *(HHEX-IDE, CDKAL1, ADCY5* and *ANK1-NXK6-3*).

**Figure 5:**
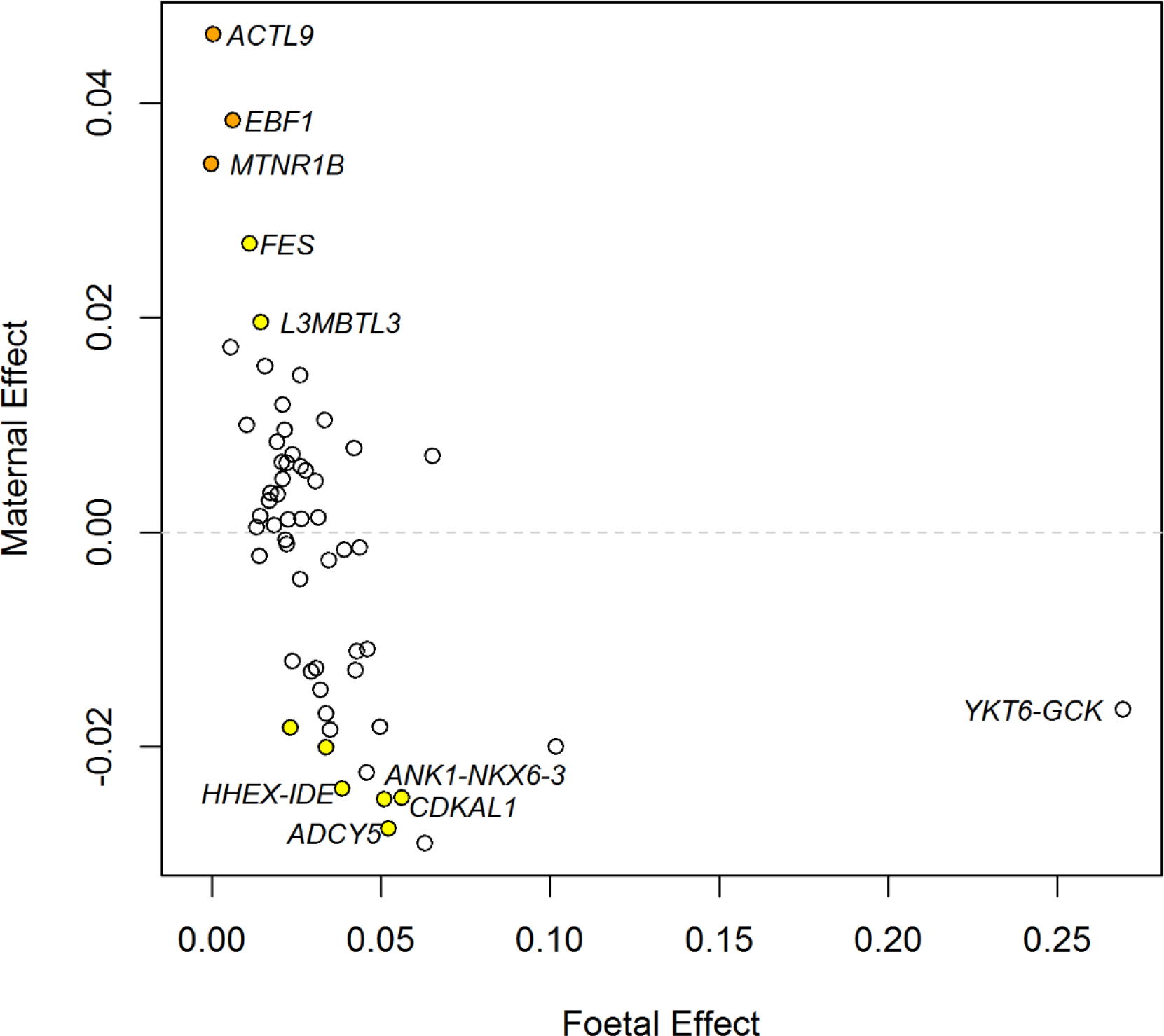
Foetal and maternal effect size estimated using the structural equation model (SEM) for the 58 birthweight associated SNPs in the UK Biobank. All SNPs are aligned to the birthweight increasing allele reported in Horikoshi *et al.*(6). The colour of each dot indicates the maternal genetic association P-value for birthweight generated using the Wald test: orange, P<0.001; yellow, 0.001≤P<0.05; white P≥0.05. Gene names are provided for those loci with large effects.

Supplementary Table 4 and Supplementary Figure 2 present the full results from the two linear models (foetal and maternal effects) and the SEM. These results show that for SNPs where the maternal and foetal effects go in opposite directions (for example the *HHEX-IDE, CDKAL1, ADCY5* and *ANK1-NKX6-3* loci), the foetal effect estimated in the GWAS(6) would have been reported to be smaller than its true effect.

## Discussion

In this article, we have presented a method for estimating and testing maternal and foetal effects. The approach uses data from mother-offspring pairs for whom genotype data are available for the mothers only and phenotype data is available on both individuals. It can also incorporate males with their own phenotype and genotype data, but not their mothers. Our method is (asymptotically) unbiased when both maternal and foetal effects exist, which improves on the traditional linear model which estimates each effect separately while assuming the other to be absent. The approach is flexible and can be used when either the individual’s own phenotype or the phenotype of their offspring is not available.

To illustrate the method, we used birthweight because there is clear evidence that both maternal and foetal effects exist(2–4, 10). However, the method could be useful for many other phenotypes, especially pregnancy outcomes and early developmental traits. As long as phenotype information is present for individuals in two generations and genotype information is available on the older individual then it is possible to use this method to estimate both maternal and fetal effects. This could include phenotypes where genome-wide association meta-analyses already exist, such as measures of size at birth including length(11) and head circumference(12), maternal phenotypes during pregnancy such as gestational weight gain(13), or developmental phenotypes during childhood such as language development(14).

The most common study design used when trying to estimate foetal and maternal effects are to have maternal/offspring pairs or parent/offspring trios, with phenotype information on the offspring and genotype information on both the parents and the offspring. These studies are then analysed using a standard linear regression model adjusting for both the maternal and offspring genotype, which is often referred to as ‘conditional analysis’. One of the benefits of the SEM we describe here is that the coefficients for the maternal and foetal effects are on the same scale as the coefficients from a conditional analysis and therefore a meta-analysis could be conducted across multiple cohorts with different study designs. Alternatively, because the model can be fit with observed covariance matrices, if the phenotypes of the mother and offspring are both standardized and the effect allele frequency is known, then the summary statistics (allele frequency, beta coefficient from the regression model and variance of the phenotype) from an unconditional analysis for either the foetal effect or the maternal effect can be incorporated into this SEM. This makes it a potentially very powerful approach as cohorts with phenotype data and genotypes from mother, child, or both, can all be incorporated. It also avoids the need to share raw data, which can be problematic for some cohorts, but still allows for all cohorts to be included in the analysis and therefore the sample size maximized.

One of the biggest advantages of this SEM is that it is robust to missing data, either for the individual’s own phenotype or the phenotype of their offspring. This is an advantage over conditional analysis, which uses only those mother/offspring pairs that have genotype data from both persons. It can even be used when no individuals have both a phenotype measured on themselves and their offspring, however the power to detect a maternal or foetal effect is reduced and a small bias is introduced to the maternal effect estimate. There are unlikely to be many studies with this study design as the majority would have a combination of individuals with complete data or missing data for their own phenotype or the phenotype of their offspring. Additionally, it is robust to measurement error involving either the individual’s own phenotype or the phenotype of their offspring. However, increasing measurement error in both phenotypes will increase the standard errors and therefore decrease statistical power, similar to the effect of measurement error on ordinary least squares regression.

The SEM using observed covariance matrices takes approximately three times longer to compute than an unconditional linear model. Therefore, there is potential for this method to be used in large genomic studies, such as genome-wide studies. A new method for fitting SEMs in genome-wide association studies in a computationally feasible fashion has recently been developed(15), which may facilitate analyses involving more complicated models like ours. We note that tests of genetic association have traditionally been performed in the fixed effects part of SEMs (i.e., the “model for the means”). In contrast, we have modelled SNP effects in the covariance part of the model which has allowed us to model latent genotypes. We have shown that within the confines of our study, accuracy of estimates of maternal and foetal effects appear to be robust to the inherent non-normality of individual level SNP data which is to be expected in the case of exogenous variables(16).

A recent study by Horikoshi et al(6) found three SNPs that were significantly associated with birthweight using the individual’s own genotype (i.e., have a ‘foetal effect’); our analyses indicate that the effects are driven by a maternal rather than a foetal effect (variants in *MTNR1B,* and near *ACTL9* and *EBF1*). The initial finding of a foetal effect appears to be due to the bias in the linear model and therefore the foetal effect size was approximately half of the maternal effect size estimated using the SEM. We also identified six SNPs where the maternal and foetal effects were in opposite directions (variants in *ADCY5, CDKAL1* and *ABCC9* and near *HHEX-IDE, ANK1-NKX6-3* and *DTL*), of which four are in regions known to be associated with type 2 diabetes.

In summary, we describe a new method for estimating unbiased maternal and foetal effects using studies where genotype data is available for only the individual and not their offspring. We have shown that the type 1 error rate of the method is appropriate, there is substantial statistical power to detect a genetic variant that has a moderate effect on the phenotype, and reasonable power to detect whether it is a foetal and/or maternal effect. We have also illustrated that this method could be useful for accurate estimation of foetal and maternal effects in large genetic studies, such as genome-wide association studies, as the computational time is not substantially larger than the standard linear model.

## Acknowledgements

We thank George Davey Smith for comments on an earlier draft of this manuscript. N.M.W. is supported by a National Health and Medical Research Council Early Career Fellowship (grant number APP1104818). D.M.E. is funded by an Australian Research Council Future Fellowship (grant number FT130101709) and an Medical Research Council programme grant (grant number MC_UU_12013/4). This research has been conducted using the UK Biobank Resource. Access to the UKBB study data was funded by University of Queensland Early Career Researcher Grant (2014002959).

## Supplementary Material

Supplementary Table 1: Bias and power to detect a significant SNP effect (P<0.05) using both the structural equation model (SEM) and a linear model with varying maternal and foetal effect sizes and minor allele frequencies. Results include simulations with complete data on all individuals and combinations of missing data for the individual’s own birthweight or the birthweight of their offspring(see Excel spreadsheet).

Supplementary Table 2: Bias and power to detect a significant SNP effect (P<0.05) when the data was simulated with measurement error using both the structural equation model (SEM) and a linear model with varying maternal and foetal effect sizes(see Excel spreadsheet).

Supplementary Table 3: Bias and power to detect a significant SNP effect (P<0.05) when the data was simulated as in the UK Biobank (i.e., only 6 discrete values for offspring birthweight) using both the structural equation model (SEM) and a linear model with varying maternal and foetal effect sizes and minor allele frequencies (see Excel spreadsheet).

Supplementary Table 4: Results from the UK Biobank for each of the 58 birthweight associated loci using a linear model (unadjusted for the effect of the other genotype) or the structural equation model (adjusted for both maternal and foetal effects). (See Excel sheet for results).

Supplementary Figure 1: Average duration (seconds) from 1,000 simulations to estimate maternal and child effects with 95% confidence intervals using an unconditional linear model, the structural equation model (SEM) with covariance matrices and the SEM with raw data. Results are presented for simulations with an allele frequency of either 0.5 or 0.99.

Supplementary Figure 2: Forest plots of the maternal and foetal genetic effects from the linear models and the structural equation model (SEM) for each of the 58 birthweight associated SNPs in the UK Biobank. “Child Unadjusted” are the results from the linear model assessing the child effect; “Child Adjusted” are the results from the structural equation model of the foetal effect allowing for maternal effects; “Maternal Unadjusted” are the results from the linear model assessing the maternal effect; “Maternal Adjusted” are the results from the structural equation model of the maternal effect allowing for foetal effects. A) is for loci that have a maternal effect (some also have a foetal effect), B) is those that are primarily driven through the foetal genotype and C) are those that don’t appear to have a strong effect from either the maternal or foetal genotype in UK Biobank. Note the different x-axis values for the *PTCH1, YKT6-GCK*and *SUZ12P1-CRLF3* loci.

## References

1. Barker DJ, Hales CN, Fall CH, Osmond C, Phipps K, Clark PM. Type 2 (non-insulin-dependent) diabetes mellitus, hypertension and hyperlipidaemia (syndrome X): relation to reduced fetal growth. Diabetologia. 1993;36(1):62-7.

2. Magnus P. Causes of variation in birth weight: a study of offspring of twins. Clinical genetics. 1984;25(1):15-24.

3. Magnus P. Further evidence for a significant effect of fetal genes on variation in birth weight. Clinical genetics. 1984;26(4):289-296.

4. Lunde A, Melve KK, Gjessing HK, Skjaerven R, Irgens LM. Genetic and environmental influences on birth weight, birth length, head circumference, and gestational age by use of population-based parent-offspring data. American journal of epidemiology. 2007;165(7):734-741.

5. Eaves LJ, Pourcain BS, Smith GD, York TP, Evans DM. Resolving the effects of maternal and offspring genotype on dyadic outcomes in genome wide complex trait analysis (“M-GCTA”). Behavior genetics. 2014;44(5):445-55.

6. Horikoshi M, Beaumont RN, Day FR, Warrington NM, Kooijman MN, Fernandez-Tajes J, et al. Genome-wide associations for birth weight and correlations with adult disease. Nature. 2016;538(7624):248-252.

7. Allen NE, Sudlow C, Peakman T, Collins R. UK biobank data: come and get it. Science translational medicine. 2014;6(224):224ed4.

8. Hattersley AT, Beards F, Ballantyne E, Appleton M, Harvey R, Ellard S. Mutations in the glucokinase gene of the fetus result in reduced birth weight. Nat Genet. 1998;19(3):268-70.

9. Pierce BL, VanderWeele TJ. The effect of non-differential measurement error on bias, precision and power in Mendelian randomization studies. International journal of epidemiology. 2012;41(5):1383-1393.

10. Ounsted M, Scott A, Ounsted C. Transmission through the female line of a mechanism constraining human fetal growth. Annals of human biology. 1986;13(2):143-151.

11. van der Valk RJ, Kreiner-Moller E, Kooijman MN, Guxens M, Stergiakouli E, Saaf A, et al. A novel common variant in DCST2 is associated with length in early life and height in adulthood. Hum Mol Genet. 2015;24(4):1155-68.

12. Taal HR, St Pourcain B, Thiering E, Das S, Mook-Kanamori DO, Warrington NM, et al. Common variants at 12q15 and 12q24 are associated with infant head circumference. Nat Genet. 2012;44(5):532-538.

13. Warrington N, Richmond R, Fenstra B, Myhre R, Gaillard R, Paternoster L, et al. Maternal and fetal genetic contribution to gestational weight gain. bioRxiv. 2017.

14. Pourcain BSt, Cents RA, Whitehouse AJ, Haworth CM, Davis OS, O’Reilly PF, et al. Common variation near ROBO2 is associated with expressive vocabulary in infancy. Nature communications. 2014;5:4831.

15. Verhulst B, Maes HH, Neale MC. GW-SEM: A Statistical Package to Conduct Genome-Wide Structural Equation Modeling. Behavior genetics. 2017;47(3):345-359.

16. Bollen KA. Structural Equations with Latent Variables: Wiley; 1989. 528 p.

